## Supplementary Information for "Eliciting priors and relaxing the single causal variant assumption in colocalisation analyses"

Chris Wallace

November 11, 2019

| Study | Data–type | $p_1, p_2$ | $p_{12}$ | Notes |
|---|---|---|---|---|
| [42] | eQTL–methQTL | empirical[1] | empirical[2] | |
| [25] | GWAS–eQTL | default | default | MS lead SNPs with eQTL signals; independent snps in each region |
| [41] | GWAS–molQTL | default | default | |
| [34] | GWAS–eQTL | default | default | |
| [23] | GWAS–methQTL | default | default | |
| [38] | GWAS–GWAS[3] | default | default | tested in 50kb windows around each of 6472 snps in HLA region |
| [22] | GWAS–chromQTL | default | empirical[4] | |
| [18] | GWAS–eQTL | default | default | post-TWAS; |
| [26] | GWAS–eQTL[5] | default | default | 45 reQTL samples only |
| [27] | GWAS–eQTL | default | default | |
| [30] | GWAS–methQTL | default | default | |
| [24] | GWAS–eQTL | | | |

*Continued on next page*

Table S1: Applied studies which used coloc in 2018

[1] set by #indep signals/#tests

[2] considered range of p12 and compared prior mean of H4 to posterior mean of H4

[3] pediatric vs adult disease

[4] The default prior on sharing (P=1105 ) was used for all primary analyses, with priors P=1104 and P=1103 evaluated separately with all other parameters unchanged.

[5] response eQTL

| Study | Data–type | $p_1, p_2$ | $p_{12}$ | Notes |
|---|---|---|---|---|
| [31] | GWAS–eQTL | | | |
| [33] | eQTL–pQTL | | | |
| [39] | GWAS–eQTL | gwas-pw[47] | gwas-pw[47] | implemented in coloc2 `https://github.com/Stahl-Lab-MSSM/coloc2` |
| [21] | GWAS–eQTL | | | |
| [29] | GWAS–eQTL | not stated | not stated | |
| [37] | GWAS–eQTL | not stated | not stated | used to confirm primary SMR analysis |
| [36] | GWAS–eQTL | default | default | |
| [19] | GWAS–eQTL | default | default | |
| [20] | GWAS–chromQTL | not stated | not stated | |
| [35] | GWAS–pQTL | default | default | |
| [40] | eQTL–pQTL | estimated[6] | p12/(p12+p1)=0.75 | used data to inform p1, p2. used conditioning and prior beliefs to inform p12 |
| [28] | GWAS–eQTL | not stated | not stated | |
| [32] | GWAS–eQTL | default | default | |

Table S1: Applied studies which used coloc in 2018. eQTL = expression QTL; mQTL = methylation QTL; pQTL = protein QTL; chromQTL = chromatin mark QTL; molQTL = selection of molecular QTLs

[6] #eqtl or #pqtl snps/number tested

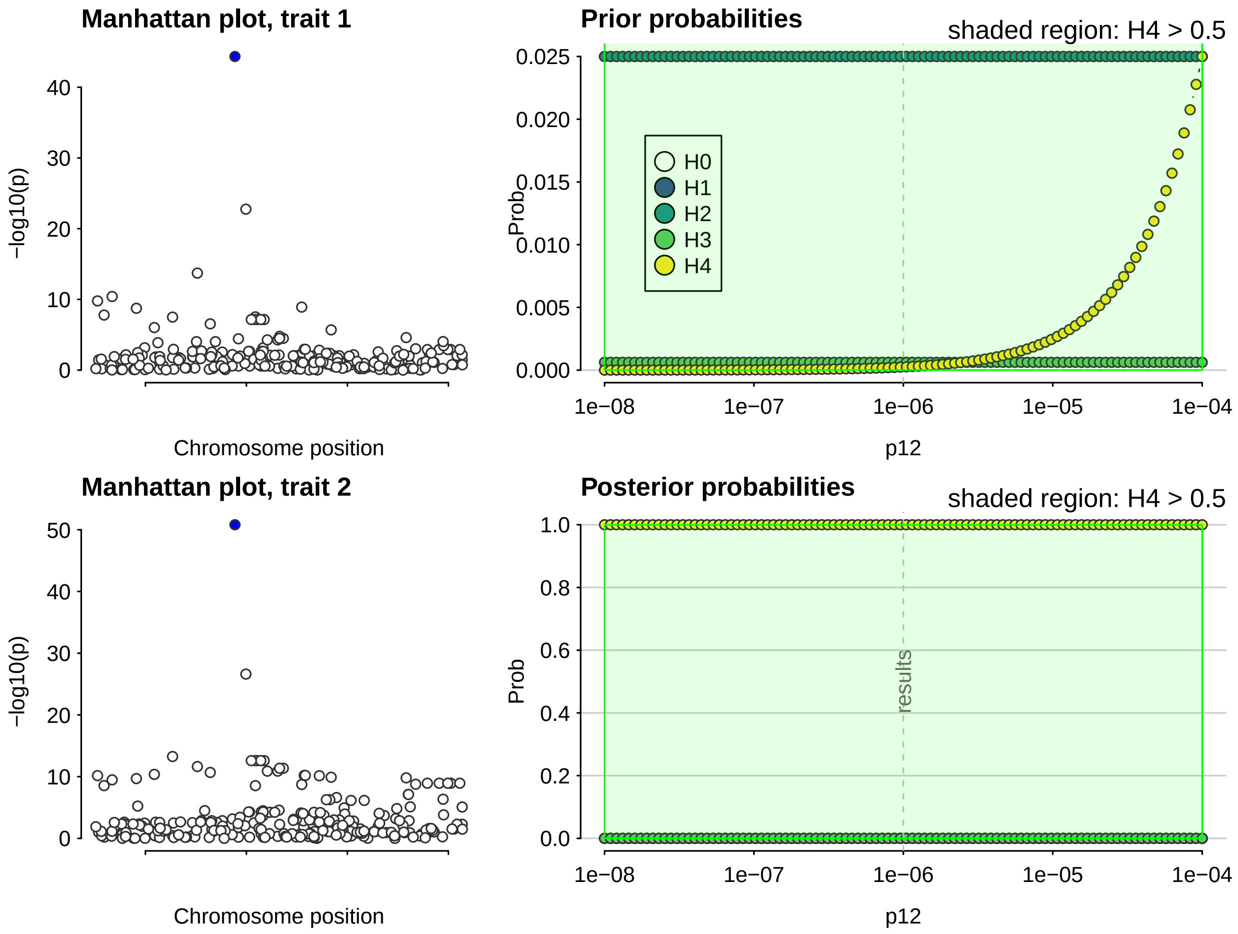


Figure S1: Example of sensitivity analysis on a dataset which shows evidence for colocalisation at a predefined rule of posterior $P(H4) > 0.5$ across a wide range of $p_{12}$. The left hand panels show local Manhattan plots for the two traits, with points coloured according the posterior probability that a SNP is causal assuming H4 is true - only one SNP has a high probability of being causal. The right hand panels show prior and posterior probabilities for H0-H4 as a function of $p_{12}$. $H_0$ is omitted from the prior plot to enable the relative difference for the other hypotheses to be seen.

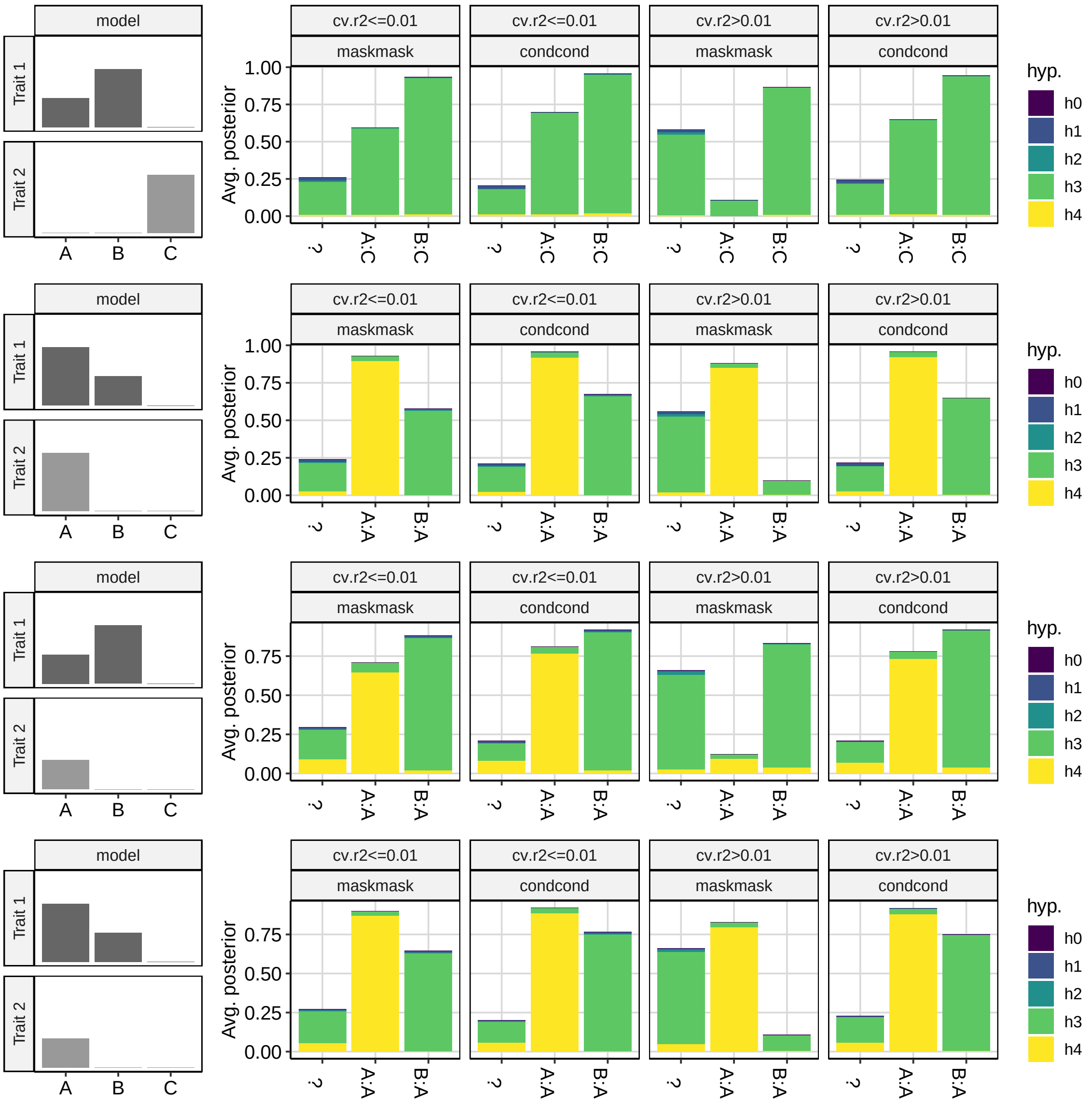


Figure S2: Average posterior probabilities for each hypothesis according to whether the maximum $r^2$ between multiple causal variants is $\leq 0.01$ or $> 0.01$. Only maskmask and condcond strategies are shown, but consistent results are seen for maskcond and condmask. Trait 1 has two causal variants, A and B, and trait 2 has just one. The left column shows the identity of causal variants for each trait and their relative effect sizes under four different models. The right column shows the average posterior that can be assigned to specific comparisons for of variants for trait 1 : trait 2. When the analysis cannot be assigned to a specific pair - i.e. when the $r^2$ between the lead SNP and the simulated causal variant is $< 0.8$, the analysis is labelled "?". Each column on the right hand side is labelled by the $r^2$ threshold used for a masked coloc analysis, and the conditional analysis is shown for reference.

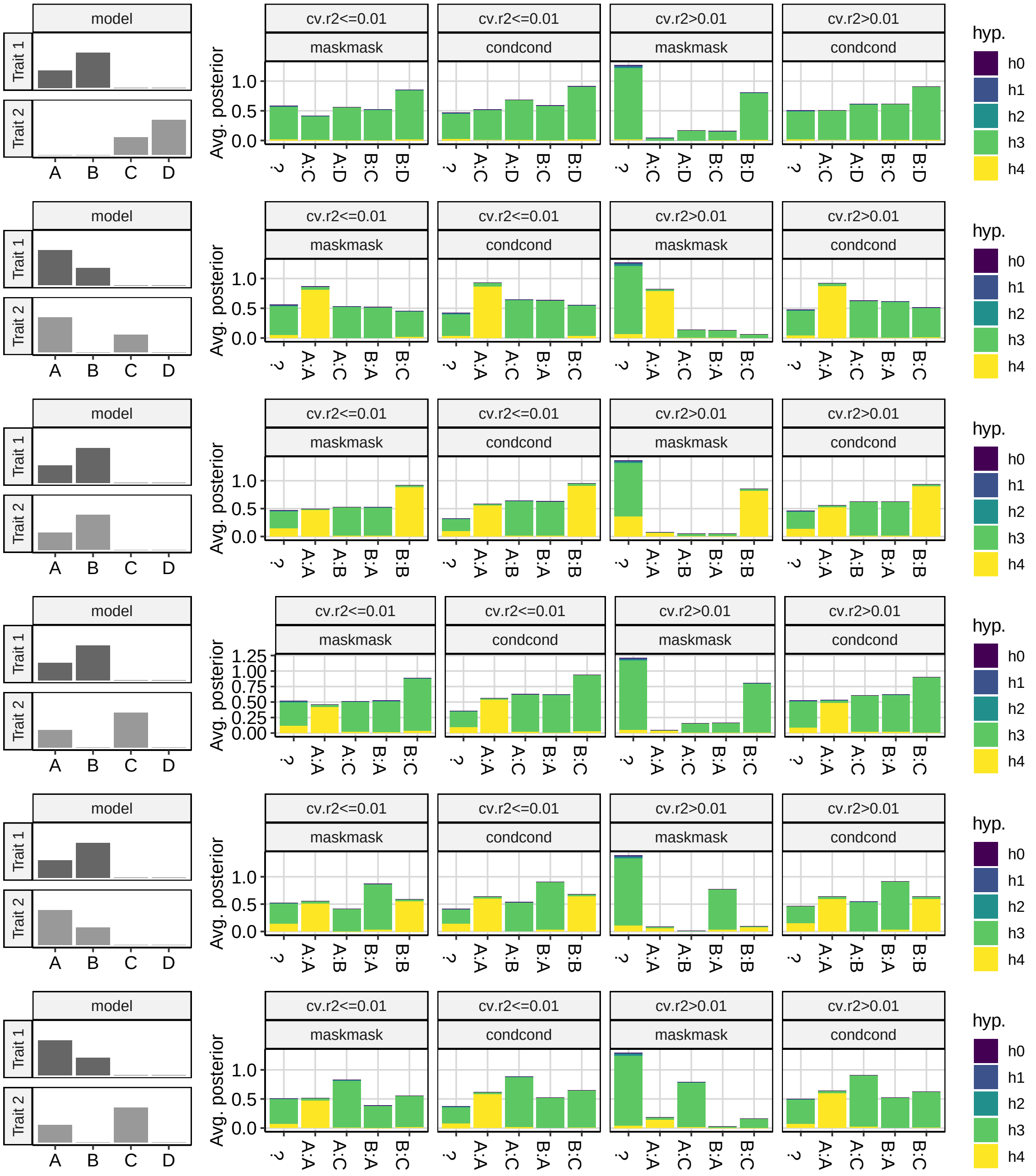


Figure S3: Average posterior probabilities for each hypothesis according to whether the maximum $r^2$ between multiple causal variants is $\leq 0.01$ or $> 0.01$. Only maskmask and condcond strategies are shown, but consistent results are seen for maskcond and condmask. The left column shows the identity of causal variants for each trait and their relative effect sizes under four different models. The right column shows the average posterior that can be assigned to specific comparisons for of variants for trait 1 : trait 2. When the analysis cannot be assigned to a specific pair - i.e. when the $r^2$ between the lead SNP and the simulated causal variant is $< 0.8$, the analysis is labelled “?”. Each column on the right hand side is labelled by the $r^2$ threshold used for a masked coloc analysis, and the conditional analysis is shown for reference.
